# Supporting the implementation of chemometrics models in bioprocess from development to commercial stage

**DOI:** 10.1101/2025.11.17.688770

**Authors:** Thibault Helleputte, Thomas Cornet, Ahmed Kanfoud, Thomas des Touches, Pascal Gerkens, Patrick Dumas, Antonio Gaetano Cardillo, Gaël de Lannoy

## Abstract

Manufacturing bioproducts involves using living organisms to produce a substance of interest. Cell culture is a crucial yet highly variable step. Monitoring, predicting, or even controlling the course of this growth requires data. For multiple reasons, the perspective of developing Process Analytical Technology based on chemometrics, rather than proceeding to actual samplings from the bioreactor, is strongly appealing and could lead to full digital twins of bioreactors. This paper addresses multiple roadblocks encountered along this modeling approach: proper model performance evaluation, limited amounts of data at process development stages, up-scaling between development and commercial stages, and regulatory concerns. These challenges are addressed by combining spectrometry data and offline readouts, simple and advanced machine learning techniques, and a good knowledge of the biomanufacturing activity and constraints. This paper uses vaccine data from multiple actual products at different scales to illustrate the considered developments.

**Authors contributions:** Thibault Helleputte, Thomas Cornet, Ahmed Kanfoud, Thomas des Touches, Pascal Gerkens and Gaël de Lannoy were involved in the conception and design of the study and/or the development of the study protocol. Pascal Gerkens, Patrick Dumas, Antonio Gaetano Cardillo and Gaël de Lannoy participated in the acquisition of data. Thibault Helleputte, Thomas Cornet, Ahmed Kanfoud and Thomas des Touches implemented the analyses, applied them to the data, and interpreted the results. All authors were involved in drafting the manuscript or revising it critically for important intellectual content. All authors had full access to the data and approved the manuscript before it was submitted by the corresponding author.

## 1. Introduction

Manufacturing biotech products such as vaccines, antigens, antibodies, and cell or gene therapies generally puts a living entity at work. These entities can be yeast, animal, or human cells, for example. This material is usually grown in a bioreactor, either for itself or to produce specific by-products (proteins, plasmids, …). This growth phase occurs typically rather upstream of the complete manufacturing process, and most of the time will strongly affect the yield of the overall process. Cell growth is a highly variable step, depending on many parameters such as feeding strategies (amount, rate, composition), temperature, pressure, growth medium, and stirring. The specific impact of the variation of these parameters (and others likely unknown) is not always well characterized. Important efforts are thus dedicated to the setup of this growth phase, during a process development phase. This phase is generally performed on a smaller scale (bioreactors of smaller volume) and in a number of experiments which is orders of magnitude smaller (typically a few dozen) than the actual number of batches that will be produced at a later commercial stage (likely hundreds or thousands). Gaining deeper knowledge about the process, such as identifying Critical Process Parameters (CPP) among Manufacturing Process Parameters (MPP), monitoring of CPP and Critical Quality Attributes (CQA) or Critical Material Attributes (CMA), is key to monitoring and controlling quality, yield and their variability. Regulatory agencies such as the US Food and Drug Administration (FDA) define Process Analytical Technology [1], or PAT, as a framework to develop, analyze, and monitor biomanufacturing processes’ quality or performance attributes taking as input timely measurements of key process parameters.

To monitor (i.e. determine in real-time or almost), predict (i.e. determine future state) or even control (i.e. adapt parameters) the course of this growth, mathematical models should be available. Such models can be mechanistic, i.e. based on extensive knowledge of actual biochemical equations describing reactions at hand [2], or hybrid, i.e. requiring a small set of data and a somehow simplified version of the biochemical equations [3], or fully data-driven. All options show benefits and drawbacks. Completely mechanistic models are highly complex (hundreds of chemical equations), most of the time even not available for a specific organism to be dealt with, and often not able to deal with the variation of baseline parameters such as pressure, temperature, etc. They represent the course of biochemical reactions given a baseline physical state. Hybrid models somehow mitigate these shortcomings but are often limited to the modeling of very few metabolites, few baseline states, etc. Yet they remain useful in the setup of feeding frequency, quantity, or composition of the initial culture medium. Purely data-driven models can address these limitations and do not require extensive knowledge of the actual reactions (black-box models). On the other hand, they will require more data, which may be difficult to obtain in large amounts at the stage of process development. Required data can be classically acquired by sampling the cell culture at repeated moments, and performing offline wet-lab analyses. The perspective of developing PAT based on spectrometry data is a much more appealing strategy, as it requires less manpower and reagents, and eliminates the risk of contamination of the culture by the sampling procedure itself. Furthermore, it increases the amount of finally available material as none is used for generating data.

The ultimate goal of such an approach is to be able, at process development time, to acquire enough spectrometry data (coupled to actual offline measurements) to build models, also sometimes referred to as Software Sensors, that could ultimately lead to complete digital twins. These models should be able to monitor the course of multiple metabolites (including the growth of the living entity of interest, but not only) with satisfactory performance. Furthermore, these models should perform well on larger-scale commercial-stage processes without requiring extensive data collection and model refitting from scratch. They must also be affordable to obtain and convincing to regulatory authorities.

This paper addresses the multiple challenges encountered in this modeling approach: proper model performance evaluation, a limited amount of data at process development stages, up-scaling between development and commercial stages, and regulatory concerns. These challenges are addressed by combining spectrometry data, offline readouts, simple and advanced machine learning (ML) techniques, and a good knowledge of the biomanufacturing activity and constraints. This paper illustrates all considered developments on actual vaccine data from multiple scales, from multiple products.

Section (2) discusses the data available for this paper and the methods used on these data. Datasets consist of a collection of Raman spectral measurements taken throughout cell culture or fermentation^1^. Each spectrum is associated with corresponding values of metabolites and biomass obtained from samples taken during spectral acquisition. This paper does not propose radically new modeling techniques but is highly original in their combination and application in this particular context. A lot of different methods are put at work in this paper, ranging from spectrometry data processing to machine learning techniques, such as feature selection [4], learning curve estimation [5], transfer learning [6] and multi-task learning [7]. Performance metrics and the protocols used for their robust estimation are also crucial aspects of the research.

Section (3) describes how these methods are combined and applied in two experimental setups to answer several pragmatic questions, such as:

- Experimental setup 1: How well does a small-scale model behave on larger-scale data?
- Experimental setup 2:

– How many development batches are required to reach satisfactory performances?
– Are there innovative functional strategies to address the limited quantity of available data at the development stage? Namely, can models be transferred from one process to another and/or be built on multiple processes simultaneously?

Each corresponding subsection describes the specific dataset used to perform each experiment and provides the corresponding results.

Section (4) discusses these results, concludes, and presents perspectives. The obtained results prove that it is possible to build and reuse (components of) models between scales and between processes. This adds credibility to the models obtained, an aspect on which utmost recent FDA draft guidance on Artificial Intelligence in GMP^2^ contexts insists [8].

## 2. Material and methods

### 2.1. Material

#### 2.1.1. Processes covered

Data from four actual vaccine products are used in this article. For confidentiality purposes, their name have been replaced by Vaccine1, Vaccine2, Vaccine3 and Vaccine4 respectively.

- **Vaccine1**

– **Process description:** A recombinant yeast strain of Saccharomyces cerevisiae was used in this work. The culture begins with an initial volume of 5.5 liters and a stirrer speed set to 260 rpm. The temperature is maintained at 30*°*C, while the pH is regulated at 5 using a base solution. The bioreactor is equipped with an in-line pO_2_ sensor that measures dissolved oxygen in percentages. The stirred motor is controlled to maintain the pO_2_ above 60%, and a peristaltic pump controls the feed flow rate. Offline measurements of optical density, glucose, acetate, ammonium, and ethanol were taken every hour. The optical density provides the concentration of biomass based on a dry weight calibration. The cultures were conducted in GSK laboratories, and further description remains confidential.
– **Reactor sizes:** 1 L or 20 L;
– **Offline measurements used:** Glucose (mM), Ethanol (mM), Acetate (mM), Ammonium (mM), Optical Density (OD at 650 nm).
- **Vaccine2:**

– **Process description:** A recombinant cell line of Chinese Hamster Ovary (CHO) cells was used in this work. The culture begins with an initial volume of 8 liters. Stirring is set to 150 rpm. Temperature is maintained at 37*°*C generally for 72 h, allowing the cells to grow, then switched to a lower value to allow them to produce the antigen. The pH is controlled at 6.95 with CO_2_ and NaOH. pO_2_ is kept constant at 30% through the addition of pure oxygen. Two solutions of nutrients are fed daily with peristaltic pumps. Offline measurements of viable and total cell densities are carried out once a day with a ViCel-XR. Filtered supernatants from the cell culture are also assayed (usually twice a day: before and after feeding) for glucose, lactate, glutamine, glutamate, ammonium, asparagine, aspartate, pyruvate, glycerol, and lactate dehydrogenase using a CedexBio-HT analyzer; and osmolarity is measured once a day. The cultures were conducted in GSK laboratories, and further description remains confidential.
– **Reactor sizes:** 4 L, 8 L, 10 L or 50 L;
– **Offline measurements used:** Glucose (g/L), Ammonium (mM), Glutamine (mM), Glutamate, Lactate (g/L), Pyruvate, total protein, LDH, Antigen (mg/L), Viable Cell Density (VCD, MCells/ml), Total Cell Density (TCD, MCells/ml). Some of these measurements have only been used in a subset of experiments.
- **Vaccine3:**

– **Process description:** A recombinant CHO cell line expressing an Vaccine3 protein was utilized, following the same culture process as used for Vaccine2.
– **Reactor sizes:** 4 L;
– **Offline measurements used:** Glucose (g/L), Ammonium (mM), Glutamine (mM), Glutamate, Lactate (g/L), Pyruvate, total protein, LDH, Antigen (mg/L), Viable Cell Density (VCD, MCells/ml), Total Cell Density (TCD, MCells/ml). Some of these measurements have only been used in a subset of experiments.
- **Vaccine4 :**

– **Process description:** Neisseria meningitidis was used in this work. A set point for pO_2_ is kept constant at 20% using a cascade of controls of stirring, air flow rate, and oxygen. Temperature was maintained at 35*°*C for the duration of the process. The pH is monitored but not controlled, in the range of 6.8*±*0.2. No feeding was performed. Samples for offline measurements are taken every 2 hours until the 8th hour and then every hour until matching the PAT experimental setup schedule. The end of the fermentation is defined based on the optical density measured with a spectrophotometer from Novaspect measuring OD at 590 nm, for which the target is 8*±*0.5. Filtered supernatants from the same OD samples were also assayed for glucose and glutamate. The cultures were conducted in GSK laboratories, and further description remains confidential.
– **Reactor sizes:** 50 L or 920 L;
– **Offline measurements used:** Glucose (g/L), Optical density (OD at 590 nm), Glutamic acid (g/L).

For Vaccine2 and Vaccine3, Raman spectral measurements were conducted on solutions of pure metabolites as well. The solutions used for the spectral acquisition included a 100 mM L-glutamic acid, 100 mM ammonium chloride, and 20 mM lactic acid.

#### 2.1.2. Spectrometry

Three spectrometry technologies have been used for generating the data used for the experiments reported in this work, depending on practical constraints not detailed here because these data have been generated in an industrial context:

- `Raman1`. Raman spectrometer from Kaiser^3^ Rxn2 with a spectral coverage of 100-3425 cm^−1^, an excitation wavelength of 785 nm and a spectral resolution of 1 cm^−1^.
- `Raman2`. Raman spectrometer from Resolution Spectra^4^ with a spectral coverage of 300-3999 cm^−1^ and a spectral resolution of 3 cm^−1^.
- `NIR`. Büchi Near-infrared spectrometer consisting of X-One V3 probe and software SX-Server, with a spectral coverage of 950-1800 nm and a spectral resolution of 10 nm.

Depending on the experimental setup, as described in Section (3), slightly different conditions apply.

### 2.2. Methods

We describe briefly in this section all methods used in the experiments covered in Section (3). All implementations have been written in R language [9].

Throughout the article, we note a measure of an endpoint for a specific batch *i* at a specific time as *y_i_*, and the measures of spectra at a given wavelength *j* for a given batch *i* at a specific time as *x_ij_*. A complete spectrum for all wavelengths is thus denoted **x***_i_*. Number of observations is noted *n*, and the number of wavelengths is noted *p*. So, for a given experimental setup, we have for each kind of endpoint (e.g. Glucose) one dataset constituted of *n* pairs (**x***_i_*,*y_i_*)^5^. Typically, the value of *n* will be the product of the number of batches available and the number of observations of the endpoint and the spectra over time (i.e. if 10 batches have been performed, and for each, 10 observations have been made for a given endpoint, then the corresponding dataset will have *n* = 100).

#### 2.2.1. Feature selection methods

Throughout the various experiments reported in Section (3), we use different methods to automatically let the model-fitting step identify a subset of the wavelengths to contribute to the model. In machine learning terms, these are called feature selection methods [4]. The following feature selection methods are used:

- LASSO [10] is a variant of the Least Angle Regression framework, which minimizes a standard mean square error loss function 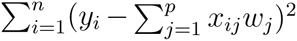 but subject to an additional regularization term which is linear a *L*1-norm over the model parameters 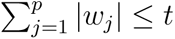 where *w*’s are the linear model parameters and *t* can be made arbitrarily small. Although initially a regression method, LASSO can be used strictly for its ability to select subsets of features, and that is how it is used in this paper.
- Random Forest [11] (referred to as “RF” in the rest of the text) is a method implementing a bagging of decision trees, for classification or regression tasks. Inside RF is a method for evaluating the overall merit of each feature. According to that metric, one can identify subsets of features of arbitrary sizes. We set this method to grow a very high (10000) number of trees in each forest to enhance feature selection stability [12].
- Partially Supervised feature selection (referred to as “PS” in the rest of the text) is a method described in [13] and which enables selecting features in the context of regression tasks while taking into account prior knowledge. The regression problem is similar to LASSO, except that (i) the regularization term is based on a *L*0-norm instead of an *L*1-norm and (ii) this regularization has a different intensity depending on prior knowledge. In this work, prior knowledge comes from spectra and offline measurements acquired on pure metabolites (sometimes diluted for practical concerns). This prior knowledge then favors the selection of some spectral wavelengths rather than others, but as a soft constraint, balanced with observation from actual data contained in datasets of interest. This method, when applicable, has been shown to increase feature selection stability while preserving or increasing predictive performances.
- Ensemble Correlation is a basic yet robust and effective feature selection method. For each given dataset, a random subsampling is performed, and Pearson correlation between each feature and the target *y* is computed. One vector of size *p* of correlations is obtained for each random subsampling. For each feature, the average correlation is calculated. A subset of features of an arbitrary size can be identified, containing the features most correlated to the target on average. This method is made available to the experimental setups of Section (3), but appears to be outperformed by one or the other of the available methods, in this case. We still mention it for the sake of completeness.

A multi-task approach is also adopted in the second experimental setting reported in Section (3.2). Multi-task, both for regression and feature selection, is described in the dedicated Subsection (2.2.3) below.

#### 2.2.2. Regression models

For the experiments reported in Section (3), we used two main regression modeling approaches. The first one is a regression version of the Random Forest (RF), and the second is a simple regularized linear model (with a two-norm regularization term, hence a ridge regression) (RLM). The rationale behind these choices is to have both a non-linear (potentially more powerful but also potentially more prone to overfitting) and a robust linear approach (less expressive, but less prone to overfitting).

#### 2.2.3. Multi-task learning

Multi-task learning is a machine learning framework for simultaneously learning multiple related regression models, on multiple datasets representing each a “task” [14]. The underlying hypothesis is that the different datasets are drawn from a common, unknown, underlying distribution [15]. In practice, the tasks have to be similar enough according to a specialist in the subject matter. Models are learned per regression task and thus per dataset, but with constraints between them. The problem is phrased as follows :

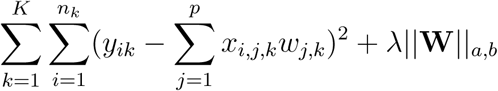

where *K* represents the number of tasks, *n_k_* the number of points in task *k*, while *x*, *y*, *w*, *i*, *j* and *p* have the same meaning as above. Uppercase and bold **W** represents the matrix of all parameters of all models (models for all tasks). *||.||_a,b_* represents various configurations of matrix norms. In this case, we use *||***W***||*_2,1_ to favor sparsity in the models, but with a shared sparsity pattern. Otherwise said, the models for all tasks will be based on a subset of the features and this subset will be common for all tasks at hand. For example, in Section (3.2) this method will select the same wavelengths in the models for the glucose over two different processes (on the same platform).

Finally, *λ* is a hyperparameter allowing to emphasize either the commonality between models (more weight on the matrix-norm regularizer) or the individual model performances (more weight on loss function). Depending on the value of this parameter, this technique can be used either just for the sake of selecting features across tasks, for multi-task regression, or both, as it is the case also for other methods like RF and LASSO. We denote both possibilities as “RMTL” throughout the text. The context indicates if it is used as a feature selection and/or as a regression method.

Multi-task learning in this work has been implemented based on the RMTL [7] package.

#### 2.2.4. Learning curve analysis

In machine learning, a learning curve [5] is a curve depicting the evolution of the performances of a model as the size of its training set increases. Even with a fixed-size dataset, this analysis can prove useful for extrapolating the amount of data that should be made available later on to reach some desired level of performance. If the curve keeps improving as the size of the training set increases, then a larger dataset size may be required (unless the performances obtained are already satisfactory). If the curve shows a plateau at an unsatisfactory performance level, then the modeling strategy should be reconsidered, rather than increasing the dataset size. In practice, for a given modeling strategy (comprising preprocessing, feature selection, and regression modeling, for example), we start by providing a small-sized training set (say 3 batches). We observe the performance of the resulting model on a test set. We repeat the same procedure except that we now provide more samples to the modeling pipeline (4, 5, 6,…), observing at each increase of training set size the impact on performances. In order to provide consistent results, the test set should be the same for all training set sizes. This whole procedure can be embedded into a resampling strategy, each time stochastically varying the composition of the test set, as explained in Section (2.2.6).

#### 2.2.5. Performance metrics

Two different aspects are reported in the experiments of Section (3). One is how well the regression models predict the concentration of the metabolites of interest compared to the ground truth as measured traditionally. The other is related to the similarity or difference of the sets of wavelengths (features) that are included or not in a given model, as a result of the feature selection step, over the multiple resampling iterations of the evaluation protocol used (see Section (2.2.6)). This second aspect is also known as the stability of the feature selection.

Two classical measures of error quantification are used to assess regression: Root Mean Square Error (RMSE) and Relative Absolute Error (RAE). A classical Pearson correlation metric is also reported.

Stability is measured via the Kuncheva index [16], a metric that compares two (or more) sets of selected features, and produces a higher score if the sets are similar. Interestingly, this score takes into account the probability of picking features at random, by taking also into account the total number of available features before selection. The larger the sets of selected features, or the smaller the total number of features, the stronger this correction.

#### 2.2.6. Evaluation protocols

Estimating the performances of predictive models is a complex topic. The classical machine learning performance estimation paradigm foresees splitting the available data in two, where a test set is left untouched^6^ until final model performances must be evaluated. On the other part of the data, resampling procedures are applied, such as cross-validation, where the data is randomly and repeatedly split in two: a train set to build various models, a validation set to estimate their performance, the combination of both enabling the tuning of hyperparameters and choosing between modeling approaches.

We follow this line of reasoning in this work while accounting for an additional specificity: data points are in our case spectra, but several spectra belong to the same production batch. To avoid obtaining too optimistic performance estimation, we impose that different spectra from a given batch be always considered in the same part of the dataset. Accordingly, all spectra from a given batch will belong either to a train set together or to a validation set together, or to a test set together. We call this protocol Leave-One-Batch-Out (LOBO).

## 3. Experiments

### 3.1. Experimental setup 1: From process development to commercial manufacturing

To address this question, we proceed as illustrated by Figure (1): The spectra and the endpoints (offline) measurements performed during cell cultures at a certain process scale are aligned. For each endpoint, a model is built using the pretreated spectra, performing feature selection and regression. The different (non-multi-task) feature selection and regression methods described in Section (2.2) are available during this training phase. The performances of these various combinations are estimated within a LOBO procedure. The most effective combination of model and feature selection for each endpoint is determined by the maximum Pearson correlation obtained. Then, the retained models are used to predict the concentration of the end-points at another scale using only the spectra obtained at this other scale. Prediction performances are calculated by comparing these predictions with the actual values of the corresponding endpoints at this other scale.

**Figure 1:**
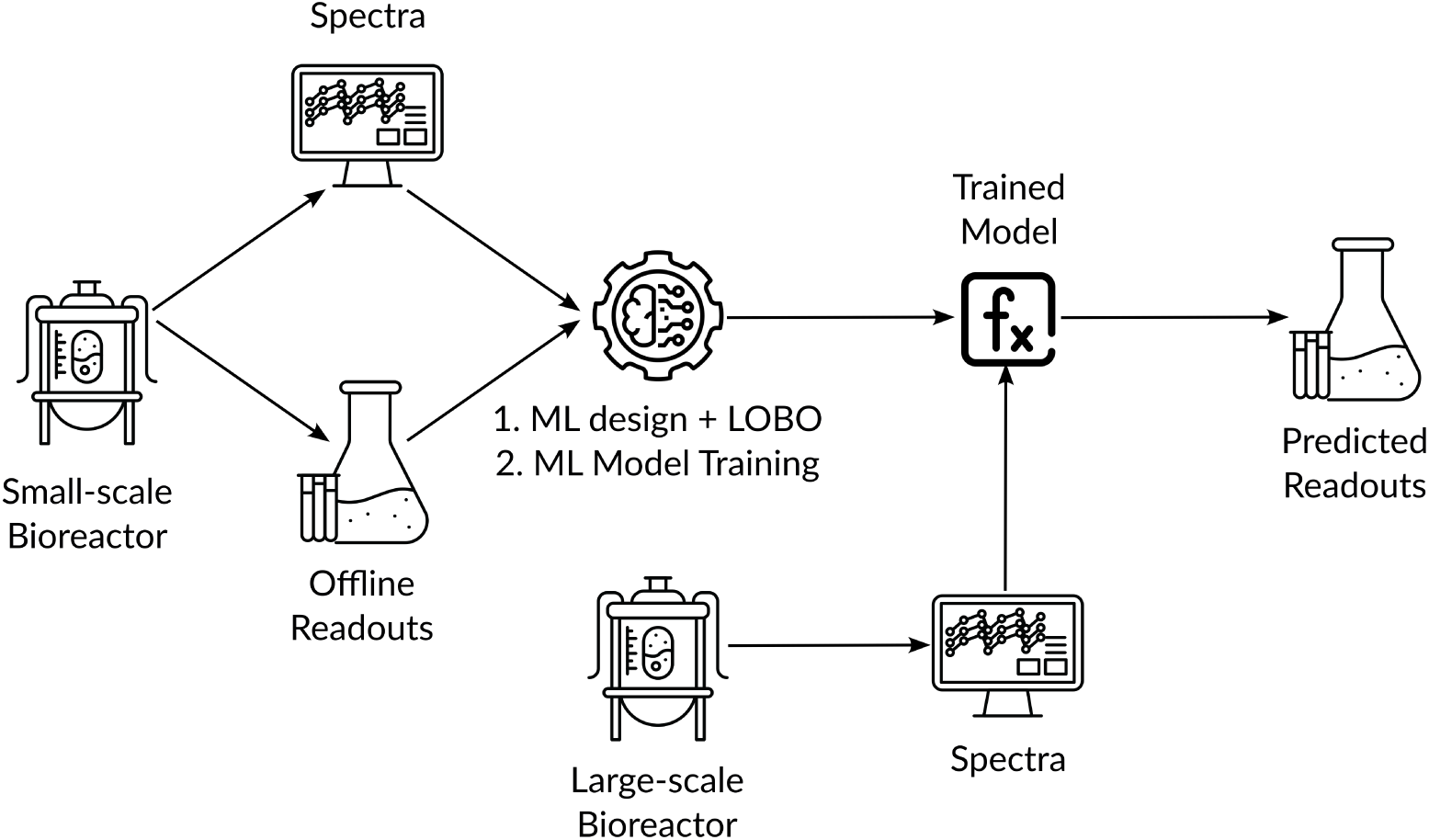
Illustration of the up-scaling experimental scheme.

#### 3.1.1. Datasets used

Table (1) gathers the datasets used in these experiments, involving three processes.

**Table 1:** Characteristics of datasets used in experimental setup 1. S stands for Small-scale, L for Large-scale.

| Project (Platform) | VACCINE2 (*CHO*) |  | VACCINE4 (*E. coli*) |  | VACCINE1 (*S. cerevisiae*) |  |
| --- | --- | --- | --- | --- | --- | --- |
| Scale (Volume) | S (4 L) | L (10 or 50 L) | S (50 L) | L (920 L) | S (1 L) | L (20 L) |
| #Batches | 13 | 3 | 10 | 3 | 3 | 13 |

For Vaccine2 process, `Raman2` spectra have been generated. Features above 3000 cm^−1^ were excluded from the analysis following GSK experts’ recommendations. Acquisition duration is 45 seconds and the number of spectra per sample is 20. Raw spectra have been detrended by 2nd derivative filter with a window size of 147 cm^−1^. Spectra are acquired every 45 minutes.

For Vaccine4 process, `NIR` spectra have been generated. Acquisition duration is 3.5 seconds. Raw spectra have been detrended by 2nd derivative filter with a window size of 490 nm. Spectra are acquired every 3.5 seconds.

For Vaccine1 process, `Raman1` spectra have been generated. Features above 3000 cm^−1^ were excluded from the analysis following GSK experts’ recommendations. Acquisition duration is 10 seconds and the number of spectra per sample is 30. Raw spectra have been detrended by 2nd derivative filter with a window size of 49 cm^−1^. Spectra are acquired every 30 minutes.

#### 3.1.2. Vaccine2 *results*

A LOBO cross-validation procedure is applied on small-scale data (spectra and endpoints) to identify, for each endpoint, the best-performing feature selection method, regression model type, and number of features to select. The criterion used to define the best combination is the Pearson correlation between the predictions and the actual values. Table (2) reports the results of that procedure for each endpoint: best-performing model type and selection method, number of features, model performances, and selection stability. A single model is subsequently built according to these best combinations on the complete small-scale dataset. These models (one by endpoint) are then used on the spectra corresponding to batches produced at a larger scale to perform pure predictions. The predictions are compared to the actual end-point values. The corresponding performance metrics for these predictions are shown in Table (3). The results are also illustrated in Figure (2).

**Table 2:** Vaccine2 models built at small-scale (first 4 columns) and their cross-validated performances at the same scale (last 4 columns). FS and #Feat. are respectively the name of the feature selection method and the number of selected features leading to the best combination.

| Endpoint | FS | Model | #Feat. | RMSE | RAE | Pearson | Kuncheva |
| --- | --- | --- | --- | --- | --- | --- | --- |
| Ammonium | LASSO | RF | 176 | 1.8±0.54 | 0.82±0.19 | 0.76±0.11 | 0.49 |
| Antigen | RF | RF | 76 | 140.12±44.1 | 0.74±0.47 | 0.97±0.03 | 0.75 |
| Glucose | RF | RF | 16 | 1.71±0.71 | 0.55±0.365 | 0.91±0.05 | 0.80 |
| Glutamine | LASSO | RLM | 202 | 1.79±0.61 | 0.76±0.22 | 0.83±0.09 | 0.54 |
| Lactate | RF | RLM | 133 | 0.57±0.35 | 0.67±0.45 | 0.9±0.08 | 0.67 |
| TCD | LASSO | RF | 468 | 4.51±1.86 | 0.71±0.29 | 0.93±0.08 | 0.44 |
| VCD | RF | RF | 819 | 5.15±1.98 | 0.87±0.42 | 0.87±0.1 | 0.64 |

**Table 3:**
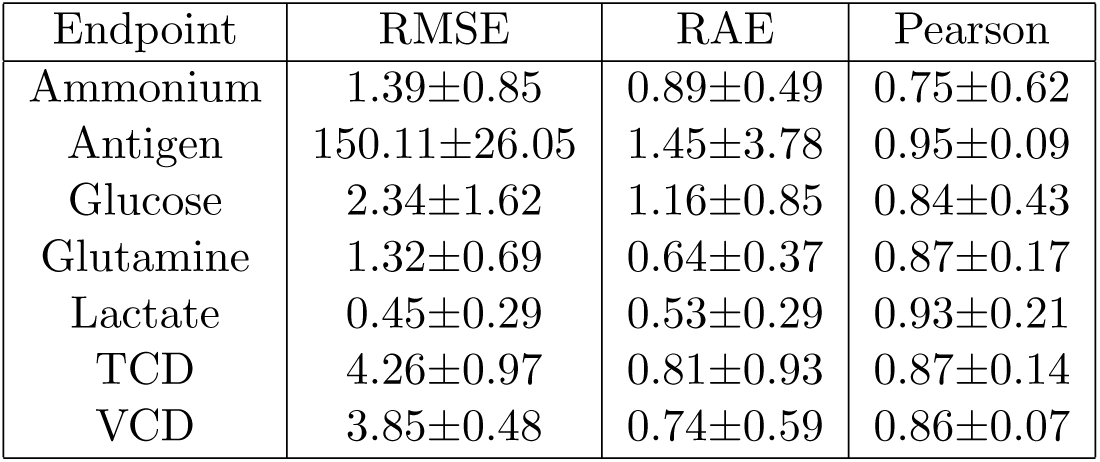
Vaccine2 predictions performances metrics on models applied on large-scale spectra when using the models trained on small-scale data.

**Figure 2:**
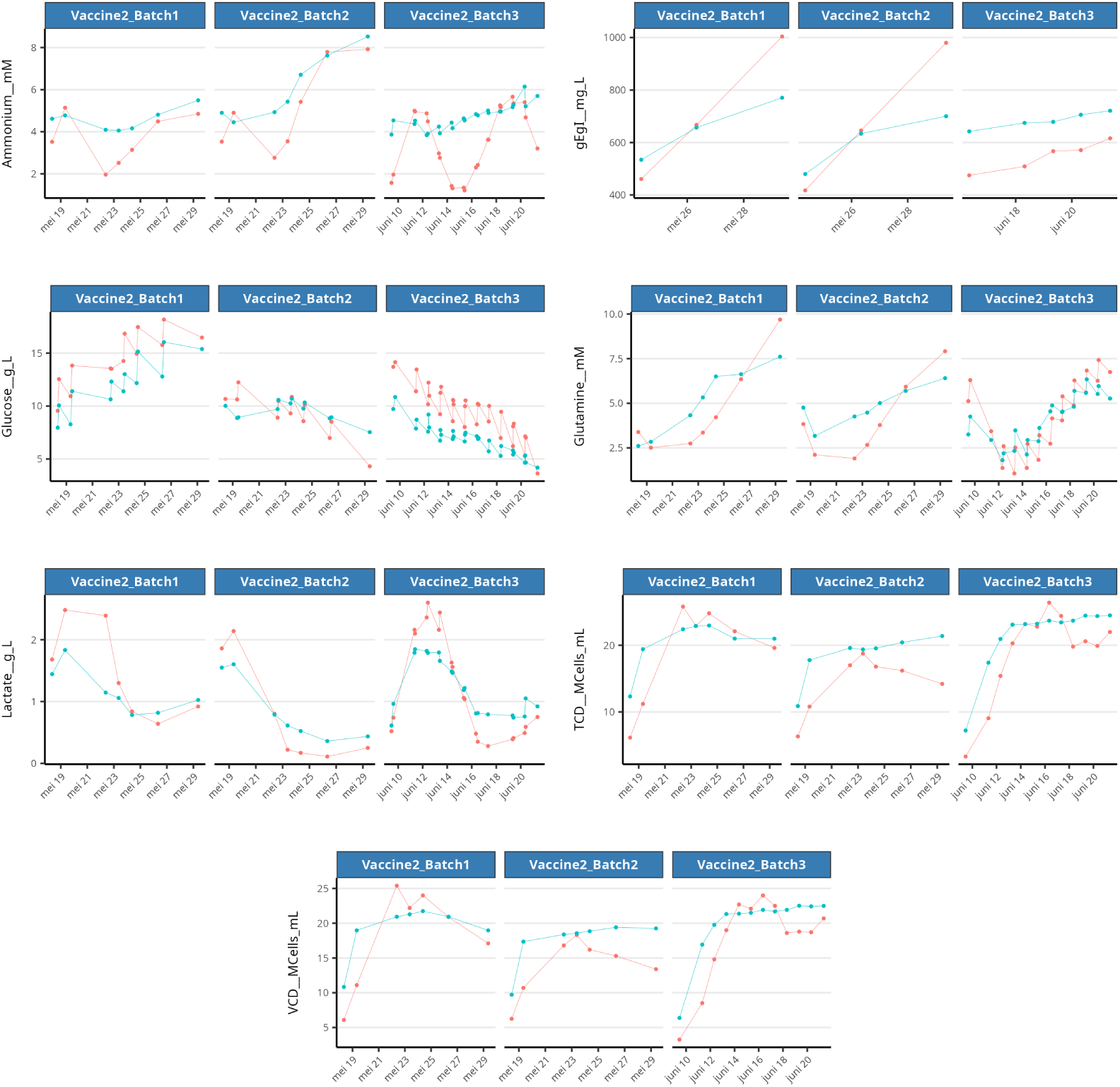
Vaccine2 predictions at large scale. In blue: the predictions of endpoints for large-scale data based on the models built on small-scale data. In red: the actual observations on the large-scale data.

#### 3.1.3. Vaccine4 *results*

The same experiment is performed for Vaccine4. Results of the LOBO procedure on the small-scale data are reported in Table (4). The models are then applied to large-scale spectra to predict endpoint values. These results are illustrated in Figure (3) and the corresponding performance metrics for these predictions are shown in Table (5).

**Table 4:** Vaccine4 models built at small-scale (first 4 columns) and their cross-validated performances at the same scale (last 4 columns). FS and #Feat. are respectively the name of the feature selection method and the number of selected features leading to the best combination.

| Endpoint | FS | Model | #Feat. | RMSE | RAE | Pearson | Kuncheva |
| --- | --- | --- | --- | --- | --- | --- | --- |
| Glucose | LASSO | RF | 16 | $0.38 \pm 0.12$ | $0.34 \pm 0.14$ | $0.99 \pm 0.01$ | 0.77 |
| OD | RF | RF | 76 | $0.24 \pm 0.06$ | $0.08 \pm 0.02$ | $1 \pm 0$ | 0.96 |
| Glutamic acid | LASSO | RF | 18 | $0.12 \pm 0.03$ | $0.11 \pm 0.03$ | $1 \pm 0.01$ | 0.95 |

**Figure 3:**
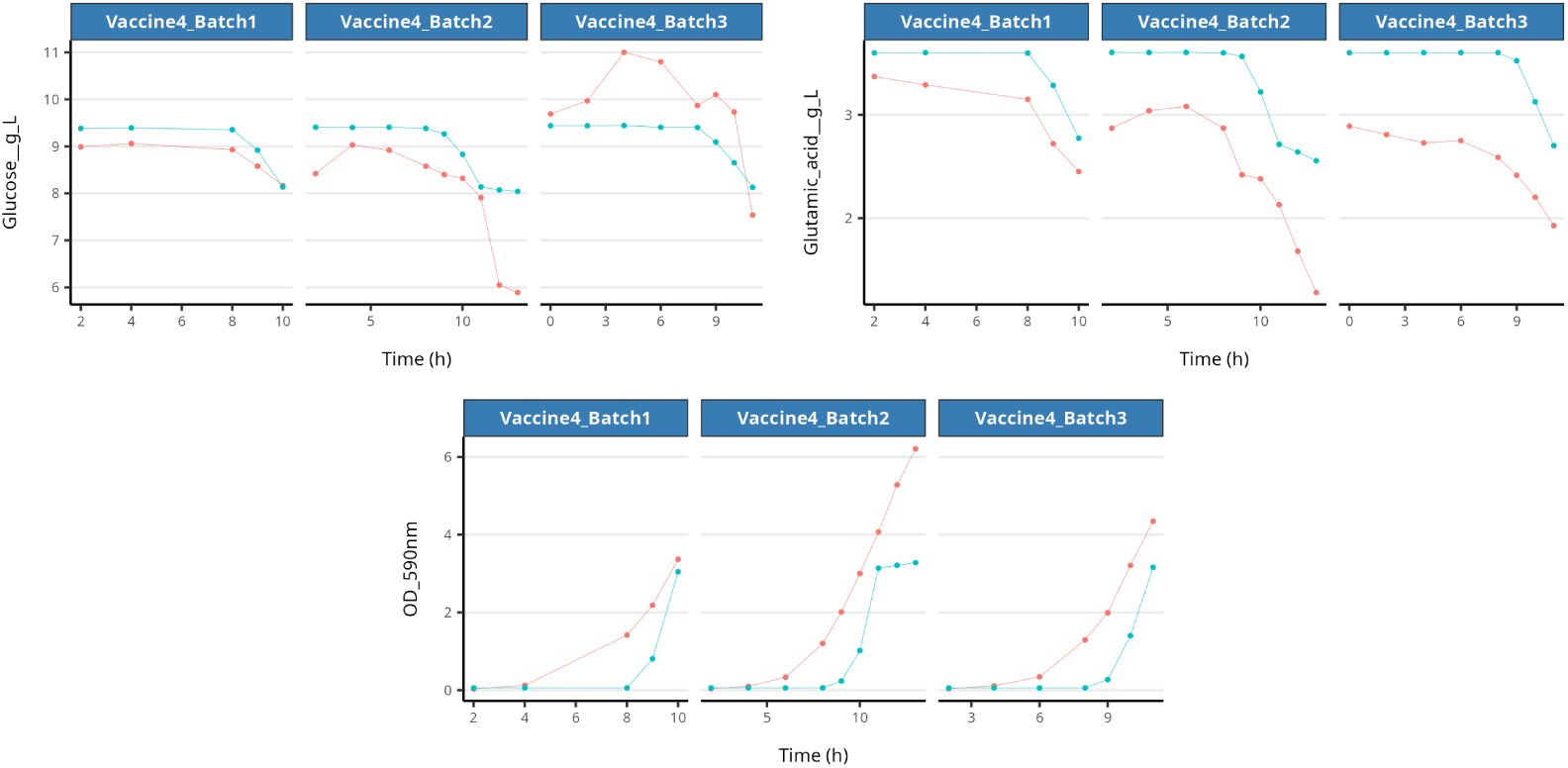
Vaccine4 predictions at large scale. In blue: the predictions of endpoints for large-scale data based on the models built on small-scale data. In red: the actual observations on the large-scale data.

**Table 5:** Vaccine4 predictions performances metrics on models applied on large-scale spectra when using the models trained on small-scale data.

| Endpoint | RMSE | RAE | Pearson |
| --- | --- | --- | --- |
| Glucose | $0.82 \pm 1.09$ | $1.13 \pm 0.59$ | $0.90 \pm 0.20$ |
| OD | $1.20 \pm 0.87$ | $0.63 \pm 0.12$ | $0.91 \pm 0.08$ |
| Glutamic acid | $0.69 \pm 0.68$ | $1.97 \pm 2.74$ | $0.93 \pm 0.03$ |

#### 3.1.4. Vaccine1 *results*

After these two successful transfers from small-scale data to a larger scale, we also seize the opportunity of having data from another process, where measurements are available in higher volumes at the larger scale rather than at the smaller scale. We thus consider the opposite exercise: transfer the models from large to small scale.

Apart from the switch of scales, everything is identical in the experimental setup as in the two previous experiments. Results are reported in Table (6), Figure (4) and Table (7).5

**Table 6:** Vaccine1 models built at large-scale (first 4 columns) and their cross-validated performances at the same scale (last 4 columns). FS and #Feat. are respectively the name of the feature selection method and the number of selected features leading to the best combination.

| Endpoint | FS | Model | #Feat. | RMSE | RAE | Pearson | Kuncheva |
| --- | --- | --- | --- | --- | --- | --- | --- |
| Glucose | RF | RF | 267 | $5.33 \pm 5.29$ | $0.19 \pm 0.16$ | $0.93 \pm 0.07$ | 0.51 |
| Ethanol | LASSO | RF | 32 | $56.87 \pm 49.57$ | $0.32 \pm 0.16$ | $0.95 \pm 0.03$ | 0.54 |
| Acetate | PS | RLM | 87 | $23.11 \pm 18.5$ | $0.5 \pm 0.18$ | $0.73 \pm 0.13$ | 0.69 |
| Ammonium | PS | RF | 133 | $92.44 \pm 18.54$ | $0.93 \pm 0.28$ | $0.5 \pm 0.15$ | 0.95 |
| OD | RF | RLM | 66 | $6.38 \pm 2.23$ | $0.15 \pm 0.05$ | $0.99 \pm 0.01$ | 0.44 |

**Figure 4:**
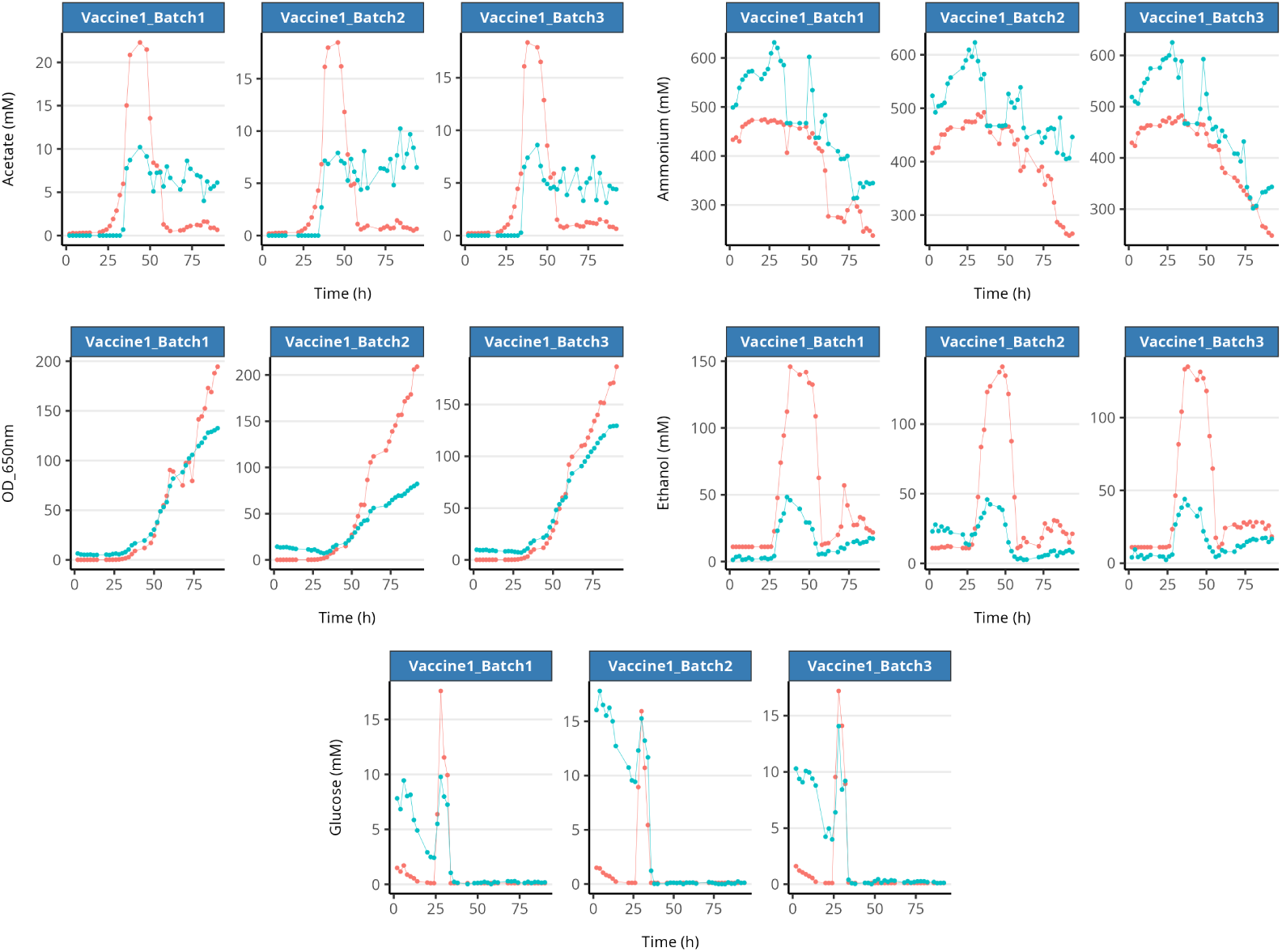
Vaccine1 predictions at small scale. In blue: the predictions of endpoints for small-scale data based on the models built on large-scale data. In red: the actual observations on the small-scale data.

**Table 7:** Vaccine1 predictions performances metrics on models applied on small-scale spectra when using the models trained on large-scale data.

| Endpoint | RMSE | RAE | Pearson |
| --- | --- | --- | --- |
| Glucose | 4.64±4.81 | 1.27±1.59 | 0.57±0.20 |
| Ethanol | 44.94±5.00 | 0.84±0.06 | 0.77±0.31 |
| Acetate | 5.16±1.21 | 0.95±0.32 | 0.43±0.30 |
| Ammonium | 94.22±27.31 | 1.25±0.57 | 0.85±0.21 |
| OD | 30.66±48.28 | 0.36±0.46 | 0.99±0.02 |

#### 3.1.4. Discussion

On all three processes and for the majority of the endpoints, the predictions of a model trained on data from a given scale and then applied on data from another scale show strikingly high correlations, despite the overall limited number of available batches for training the models. This result promotes using models at a target scale very early, benefitting from data and models from other scales. This model transfer appears to work both ways (small to large, large to small). The coherence between scales (the fact that a scale-specific model can be reused at another scale) promotes trust in these models.

In some cases, however, the predictions at the target scale are globally shifted compared to actual data, i.e. curves are somehow parallel, but lower or higher. This vertical offset impacts predictive performances in terms of RMSE or RAE, but not in terms of correlation. This might be related to the fact that correlation alone has been used as a criterion of choice in this experimental campaign to choose between combinations of model type and feature selection method at the end of the LOBO procedure.

How could this be overcome in future work? First, the models learned on a first scale could be re-calibrated to compensate for this offset on the target scale (all parameters fixed, excepted their offset that would be recomputed). This could be done using the spectra of a small number of batches at the targeted scale. An alternative option could be to measure the endpoint at a few time points on each batch at the targeted scale to re-align the corresponding predictive curve at the start of each batch.

Another investigation track could be the design of a composite score for the choice of model type and feature selection, taking into account both kinds of metrics: one based on the intensity of offset (such as RAE or RMSE) and one based on correlation, for example via a geometric mean of both.

Additionally, this composite score could also account for the stability of the feature selection. This could lead to an improvement of the lower stability results observed for example on Ammonium, Glutamine, and TCD in the experiment of Vaccine2. Evaluating such an option consisting of a composite score might not lead only to favorable outcomes: stability and correlation could be only moderate for more endpoints, for example.

### 3.2. Experimental setup 2: Overcome lack of large datasets

In this second experimental part, we address another situation encountered in some biomanufacturing companies: can we benefit from the fact that the company develops or manufactures different products on similar platforms? For example, we will use a real case where two products are based on CHO cultures: Vaccine2 and Vaccine3. These products have differences but still are based on the same living organism. Is it possible to exploit this commonality? Are there functional strategies to address the limited data available at the development stage? Namely, can models be transferred from one process to another and/or be built on multiple processes concurrently? In particular, we reproduce the following scenario: let us suppose a first process has been developed, that will be named “source task” and for which a series of data points, each composed of a spectrum and an offline endpoint measurement (**x***_i_*,*y_i_*), are already available, and that a second process based on the same expression platform has now to be developed. This second process will be named “target task”. In the experiments reported below:

- Source task includes up to 22 Vaccine3 small-scale batches.
- Target task includes up to 8 Vaccine2 small-scale batches.

The objective of this experiment is double : (i) to evaluate how the information contained in the source task (older process) may benefit the target task (newer process), and (ii) in such a case, how much data from the target task are needed to reach satisfactory performances. To address these two objectives in one experiment, we combine two techniques: multi-task learning and a learning curve, both described in Section (2.2).

Within a LOBO^7^ procedure, we adopt a multi-task approach, where multi-task models are built on both processes, but on top of that, there is an additional loop for implementing a learning curve just for the target task. So at each iteration of the LOBO, a multi-task approach is run with a fixed number of batches from the source task (22 Vaccine3 batches), but a growing number of target task training batches (Vaccine2 batches, from 1 to 7, plus 1 as a test batch). Figure (5) illustrates this experimental setup, combining multi-task learning and the evaluation of a learning curve.

**Figure 5:**
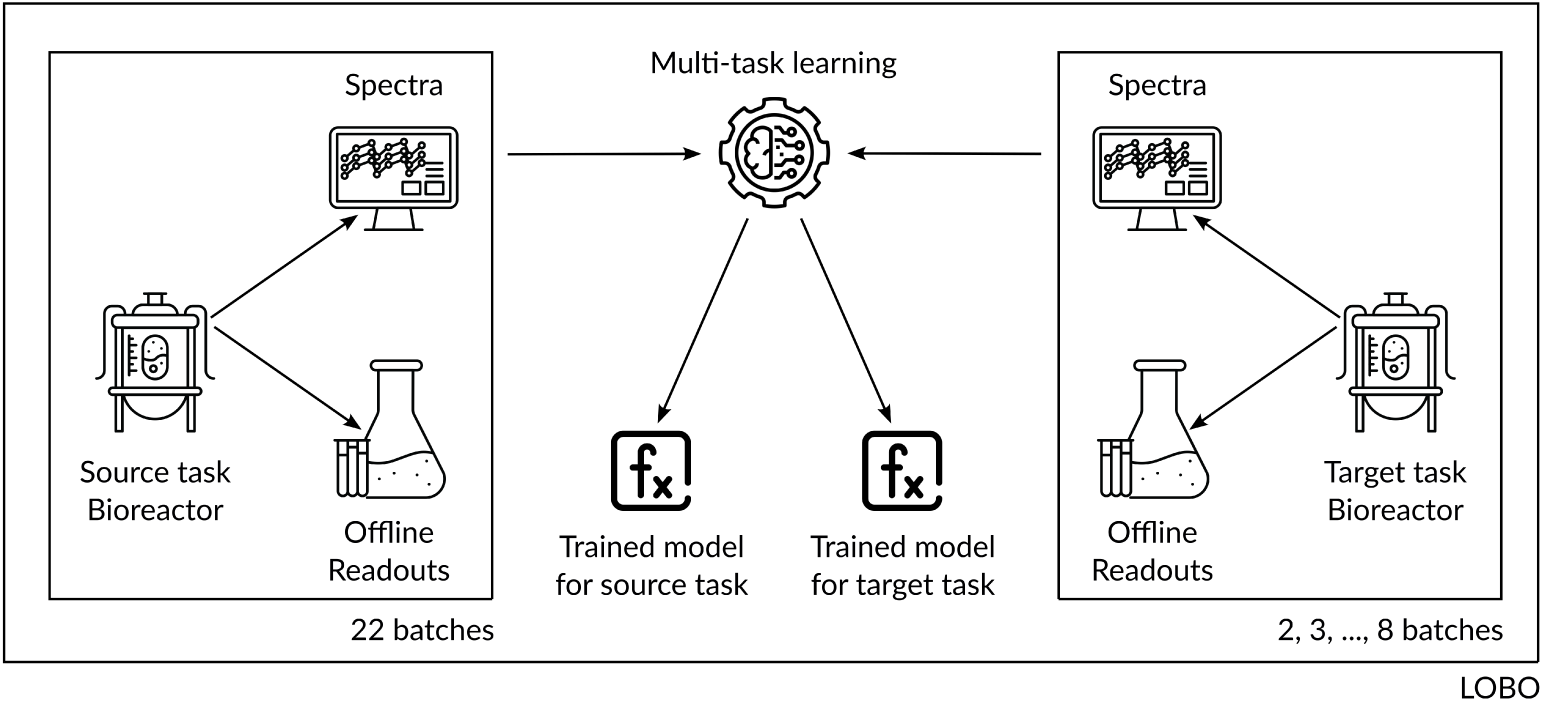
Multi-tasking and learning curve. The number of batches available for the source task is larger and constant. The number of batches for the target task is increasing and in any case lower. It starts at 2, in order to have at least one batch for training and one batch for evaluation in the LOBO loop.

The model type and number of features have been dictated by the previously developed Vaccine3 model. Two specific feature selection methods were applied to the model for each endpoint :

- PS: Applied to endpoints with available pure spectra (e.g., glucose, lactate)^8^
- RMTL: Used for endpoints without pure spectra (e.g., antigen)

The most effective model for each training size was determined using the Pearson correlation alone.

Applying this multi-task learning approach requires that all learning tasks considered contain (exactly) the same endpoints (features). This experiment focuses on four endpoints available in both datasets and the most crucial to monitor from an industrial perspective: Glucose, Glutamine, Lactate, and Antigen. Among these, antigen was treated as a control due to its anticipated lower transferability. Indeed, the two products are based on different antigens, and it is expected that their production differs, even when the cell growth itself would be similar for both products. Consequently, we hypothesized that the learning curve for antigen would show slower improvement, with poorer initial performance and requiring a larger amount of Vaccine2 data to reach acceptable predictive accuracy. This assumption provided a baseline for evaluating the model’s transferability for the other endpoints.

To extend our comprehension of the benefit of the information flow from source to target task, a second version of this experiment is conducted where the amount of Vaccine3 data in the source task is reduced from 22 batches to only 3.

Overall, the expected results are as follows:

- Performances on target task should be reasonable even with a very low number of target task batches, given the information flowing from the source task;
- Performances on target task should increase as the number of target batches increases;
- Performances on the target task should be degraded when fewer batches are available in the source task.

These expectations are checked in Section (3.2.2).

#### 3.2.1. Datasets used

Table (8) gathers the datasets used in this experiment, involving two different processes. For both Vaccine2 and Vaccine3 processes, `Raman1` spectra have been generated at small scale. Features between 3001 and 3425 *cm*^−1^ were excluded from the analysis following GSK experts’ recommandations. Acquisition duration is 45s for Vaccine2 and 30s for Vaccine3 and number of spectra per sample is 20 for Vaccine2, and 20 for Vaccine3 as well. Raw spectra have been detrended by 2nd derivative filter with a window size of 49 *cm*^−1^.

**Table 8:** Characteristics of datasets used in experimental setup 2. S stands for Small-scale.

| Project (Platform) | VACCINE2 (*CHO*) | VACCINE3 (*CHO*) |
| --- | --- | --- |
| Scale (Volume) | S (4 L or 8 L) | S (4 L) |
| #Batches | 8 | 22 |

#### 3.2.2. Results

Table (9) summarizes the model and features used for each endpoint. However, the hyperparameters, including the number of selected features and the specific features themselves, are not fixed. For each new model created, corresponding to varying training set sizes (given the learning curve context), a new set of features was selected to optimize performances.

**Table 9:**
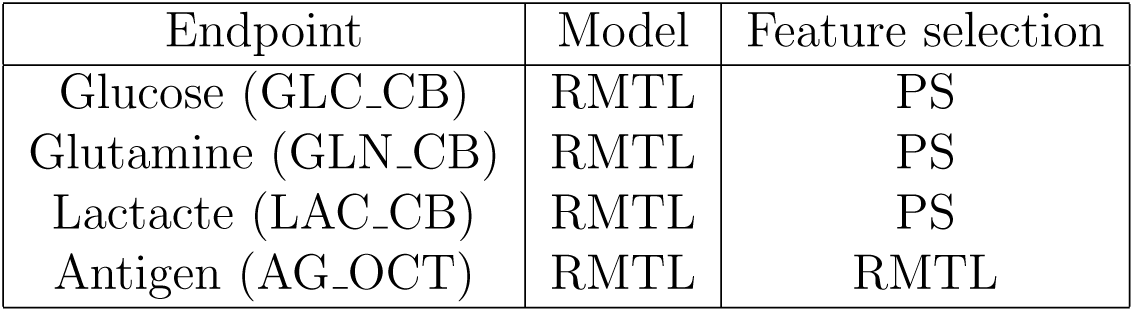
Model and feature selections used for the four primary endpoints.

Results are reported in Figures (6) to (9). The graphs for RAE, RMSE, Pearson correlation, and Kuncheva represent the mean values across all LOBO iterations for each endpoint. The shaded ribbon in each graph corresponds to the interquartile range (25%-75%). These graphs display the learning curves, i.e. how performances are impacted by the training size increase on the target task (Vaccine2). Both conditions on the amount of data available from the previous process, the source task (Vaccine3), are depicted on the same figures in a different color.

**Figure 6:**
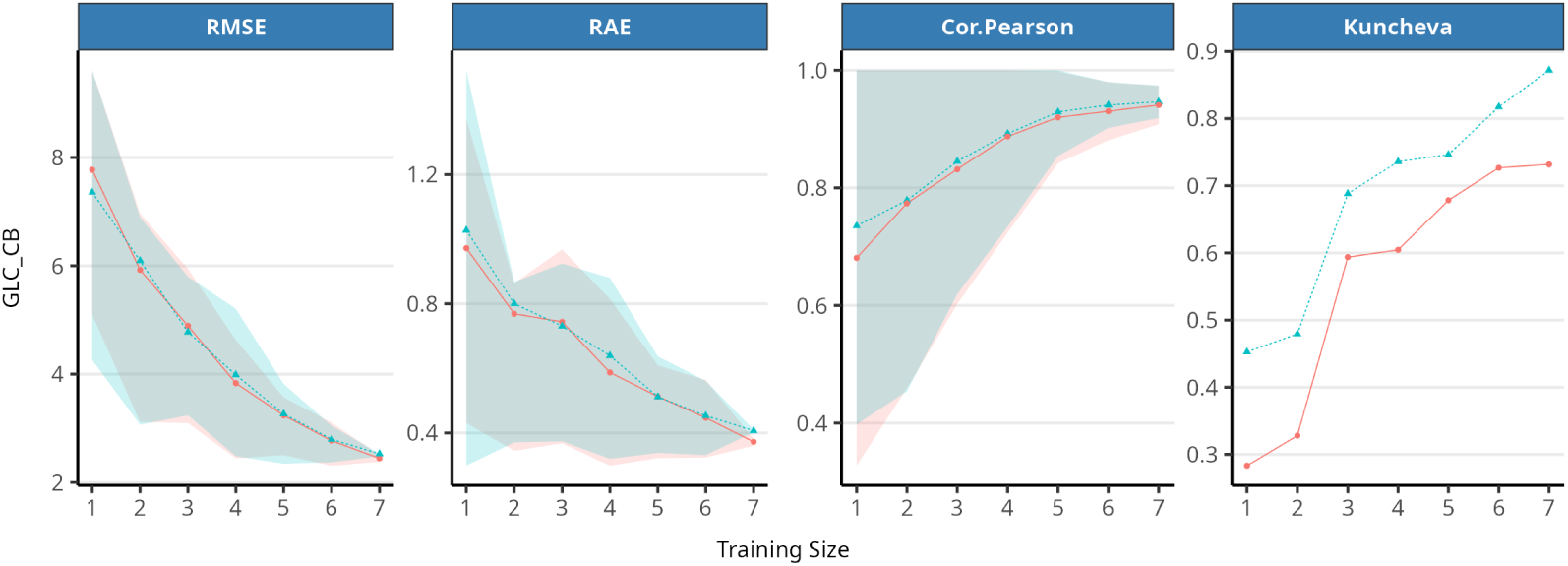
Glucose. In blue: 22 batches are included in source task (Vaccine3). In red: source task (Vaccine3) is limited to 3 batches.

**Figure 7:**
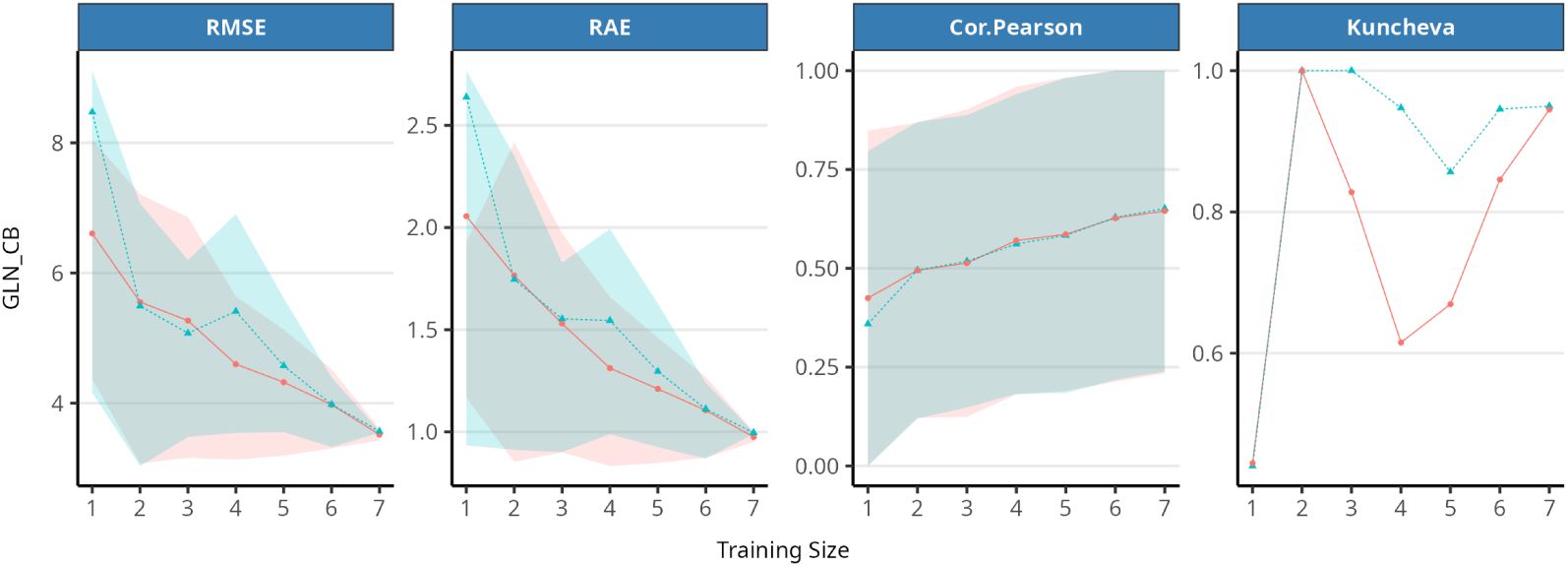
Glutamine. In blue: 22 batches are included in source task (Vaccine3). In red: source task (Vaccine3) is limited to 3 batches.

**Figure 8:**
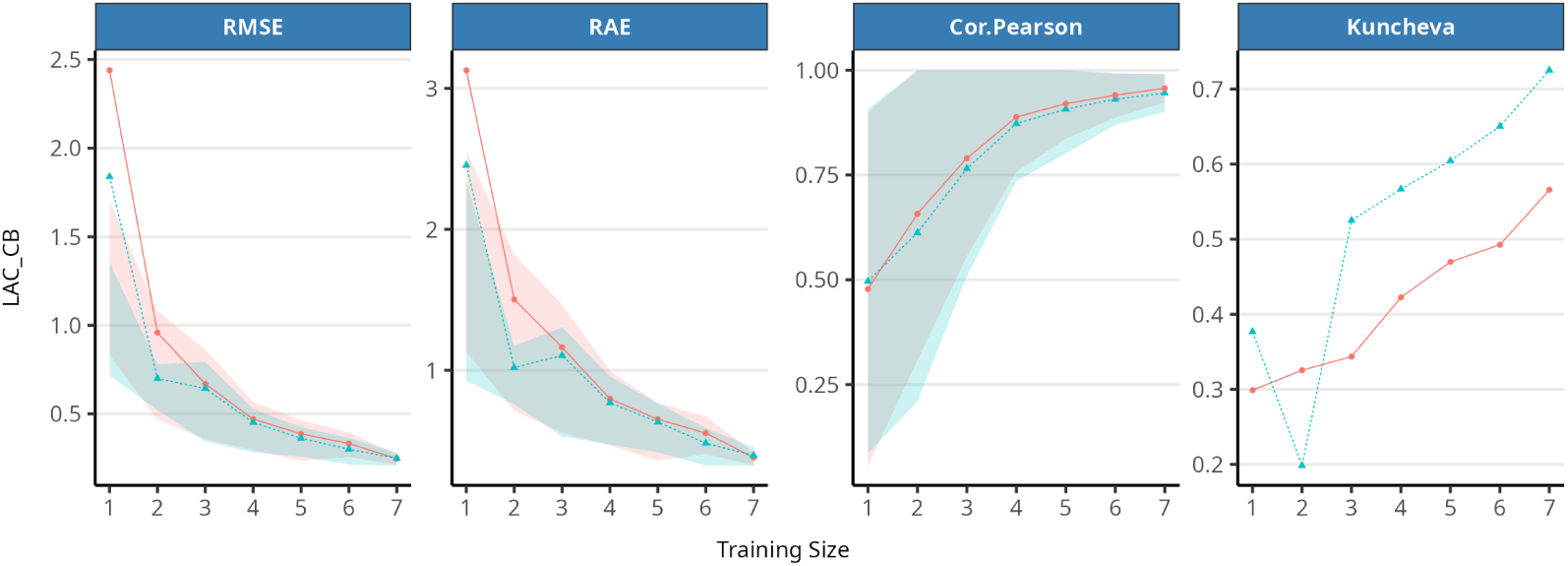
Lactate. In blue: 22 batches are included in source task (Vaccine3). In red: source task (Vaccine3) is limited to 3 batches.

**Figure 9:**
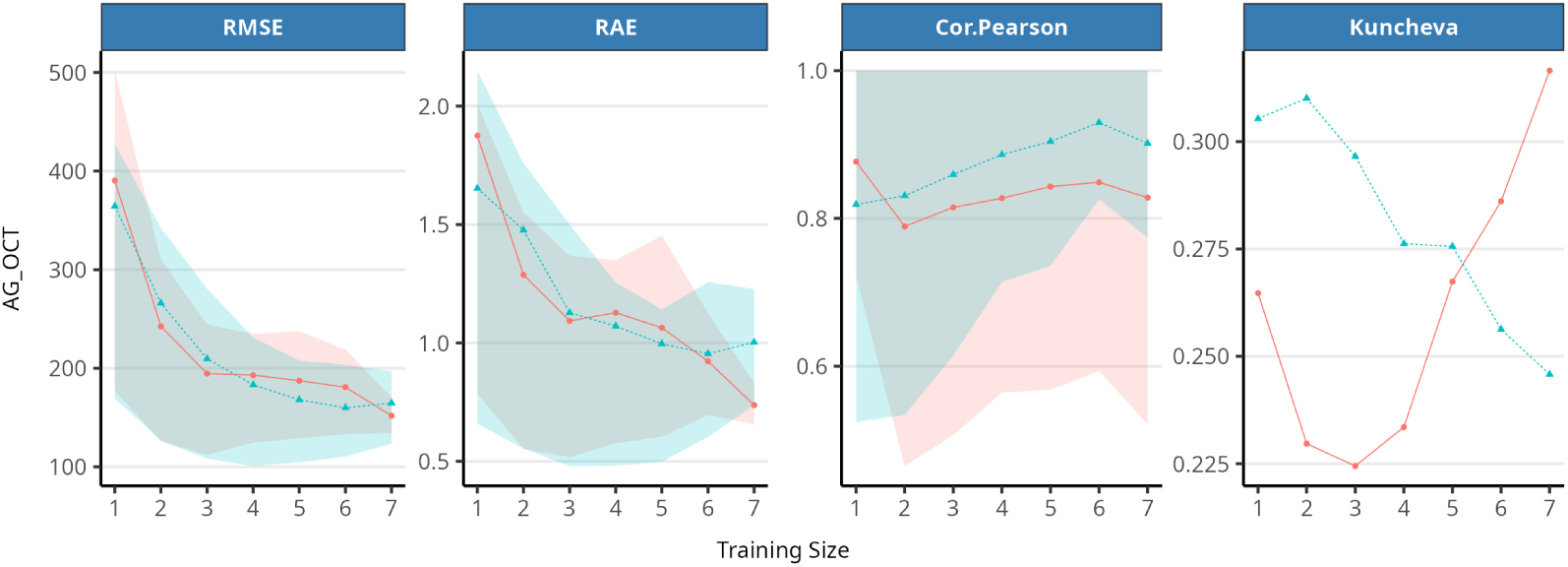
Antigen. In blue: 22 batches are included in source task (Vaccine3). In red: source task (Vaccine3) is limited to 3 batches.

#### 3.2.3. Discussion

These results show that increasing the training set size for the new process in development, the target task (Vaccine2), leads to a notable improvement in model performance. However, the extent of this improvement varies across metabolites and the evaluated metrics. For most endpoints, the learning curves show rapid progress up to approximately 4 batches, where a small plateau is witnessed for key metrics such as Pearson correlation, RAE, and RMSE. This indicates that, beyond this threshold, additional gains in average accuracy become less significant. Yet, the variability continues to drop beyond this point. Using batches from a source task allows us indeed to rapidly reach satisfactory performances for modeling a new process.

In contrast, model stability, as measured by the Kuncheva index, continues to improve significantly as the number of batches increases, suggesting that the models can maintain more consistent feature sets across resampling iterations when additional data is available.

The comparative analysis between models trained with a small number (3) of source task batches (Vaccine3) and those trained with the full set of source task batches (22) highlights a mixed impact: while increasing the batches in source task considerably improves stability (Kuncheva index), it has minimal effect on other metrics. These results indicate that additional information from the source process primarily increases consistency rather than predictive accuracy. This is another positive conclusion, as using data from several processes (different products but similar platforms) in a multitask learning setup favors the consistency of selected features (wavelengths). This should give more confidence in the models.

Finally, the distinct behavior of the antigens used as a control is a sanity check that validates the initial hypothesis that this metabolite would be challenging to transfer. Even with 7 Vaccine2 batches in the training sets, performance remains poor, with high RAE and RMSE values and low stability (Kuncheva index ranging from 0.22 to 0.35). Notably, increasing the number of batches in the source task even reduces stability for this endpoint on the target task rather than improving it as it did for the other endpoints.

On top of the four considered endpoints, additional ones with lower industrial priority such as Glutamate, Pyruvate, VCD, TCT,… have been analyzed to ensure the robustness and generality of the conclusions. The results for the majority of these extra endpoints converged toward similar observations.

## 4. Conclusions

This article shows how to practically harness spectrometry and offline data to implement chemometrics modeling during bioprocess development. In particular, this paper shows what methods can be used to build powerful models on limited amounts of data, automatically selecting useful features (wavelengths). It describes proper approaches for assessing predictive performances. Finally, it demonstrates how to implement such strategies in practice, bridging process development with commercial-stage settings, where models from a given reactor scale contribute to modeling at other scales. Applications are reported on four actual vaccine processes at various scales, with predictive performances in most cases above 90% correlation between predictions and actual values. A benefit of one of the proposed strategies (multi-task learning) is that, in total, models are built on more batches and possibly on several processes at once. This element increases the stability of the parameter selection (feature selection) and is likely to reassure regulatory agencies regarding the soundness of these models. Overall, similarities between models across scales or across processes contribute to the credibility of the models, a criterion put forward in the very recent draft guidance for the use of AI models in GMP contexts by the FDA [8].

This work demonstrates that for scaling up (or down) or when developing a new process on an existing platform, acceptable results are already obtained with spectrometry and offline data from as few as a dozen batches.

Some limitations have been identified, for which perspectives of extension of this work are proposed. In particular, one experimental choice of this work is to select models during cross-validation steps and hyperparameter tuning based on a correlation measure. As a result, correlation levels when finally evaluating model performances are very high. One may desire to favor other metrics, such as RMSE, RAE, or stability. In such a case, these measures should replace correlation during the model optimization/selection steps, or a hybrid criterion should be conceived. The result of this multicriteria approach could be evaluated in future work.

## Footnotes

1 We use both terms interchangeably in this paper, even if fermentation may sometimes be a slight abuse of language.

2 Good Manufacturing Practices.

3 Now Endress + Hauser.

4 Now Procellics from Merck.

5 It is assumed that for a pair (**x***_i_*,*y_i_*), **x***_i_* and *y_i_* are observed at the same time *t*, but *t* is omitted to simplify notations.

6 Including untouched by feature selection.

7 In this multi-task context, one batch *per task* is left out at each iteration.

8 PS is a classical, mono-task feature selection method. Hence, it applies to each task separately. It will thus produce a selected feature list or ranking for each task. One has to unify the per-task lists of selected features, for RMTL requires the same features for all tasks. This unification is achieved by an ad-hoc algorithm based on the rankings of feature importance per task. The general idea is that if a feature is important (i.e. selected or highly ranked) for one task, it will be included in the unified selection even if it is less important for the other task(s).

## References

[1] U.S. Department of Health and Human Services, Food and Drug Administration, Center for Drug Evaluation and Research (CDER), Center for Veterinary Medicine (CVM), Office of Regulatory Affairs (ORA), Guidance for Industry. PAT — A Framework for Innovative Pharmaceutical Development, Manufacturing, and Quality Assurance, Pharmaceuticals CGMPs (2004).

[2] P. Kroll, A. Hofer, S. Ulonska, J. Kager, C. Herwig, Model-based methods in the biopharmaceutical process lifecycle, Pharmaceutical Research 34 (2017) 2596–2613. doi:10.1007/s11095-017-2308-y. URL 10.1007/s11095-017-2308-y

[3] L. Dewasme, M. Mäkinen, V. Chotteau, Practical data-driven modeling and robust predictive control of mammalian cell fedbatch process, Computers & Chemical Engineering 171 (2023). 10.1016/j.compchemeng.2023.108164.

[4] I. Guyon, A. Elisseef, An introduction to variable and feature selection, Journal of Machine Learning Research 3 (2003) 1157–1182.

[5] T. Viering, M. Loog, The shape of learning curves: A review, IEEE Transactions on Pattern Analysis and Machine Intelligence 45 (6) (2023) 7799–7819. doi:10.1109/TPAMI.2022.3220744.

[6] T. Helleputte, P. Dupont, Feature selection by transfer learning with linear regularized models, in: W. Buntine, M. Grobelnik, D. Mladenić, J. Shawe-Taylor (Eds.), Machine Learning and Knowledge Discovery in Databases, Springer Berlin Heidelberg, Berlin, Heidelberg, 2009, pp. 533–547.

[7] H. Cao, J. Zhou, E. Schwarz, RMTL: an R library for multi-task learning, Bioinformatics 35 (10) (2018) 1797–1798. arXiv:https://academic.oup.com/bioinformatics/article-pdf/35/10/1797/48970181/bioinformatics_35_10_1797.pdf, doi:10.1093/bioinformatics/bty831. URL 10.1093/bioinformatics/bty831

[8] U.S. Department of Health and Human Services, Food and Drug Administration, Center for Drug Evaluation and Research (CDER), Center for Biologics Evaluation and Research (CBER), Center for Devices and Radiological Health (CDRH), Center for Veterinary Medicine (CVM), Oncology Center of Excellence (OCE), Office of Combination Products (OCP), Office of Inspections and Investigations (OII), Considerations for the Use of Artificial Intelligence to Support Regulatory Decision-Making for Drug and Biological Products. Draft Guidance for Industry and Other Interested Parties., Pharmaceuticals CGMPs (2025).

[9] R Core Team, R: A Language and Environment for Statistical Computing, R Foundation for Statistical Computing, Vienna, Austria (2021). URL https://www.R-project.org/

[10] B. Efron, T. Hastie, I. Johnstone, R. Tibshirani, Least angle regression, The Annals of Statistics 32 (2) (2004) 407 –499. doi:10.1214/009053604000000067. URL 10.1214/009053604000000067

[11] L. Breiman, Random forests, Machine Learning 45.

[12] J. Paul, P. Dupont, Inferring statistically significant features from random forests, Neurocomputing 150 (2015) 471–480, special Issue on Information Processing and Machine Learning for Applications of Engineering Solving Complex Machine Learning Problems with Ensemble Methods Visual Analytics using Multidimensional Projections. 10.1016/j.neucom.2014.07.067. URL https://www.sciencedirect.com/science/article/pii/S092523121401234X

[13] T. des Touches, M. Munda, T. Cornet, P. Gerkens, T. Hellepute, Feature selection with prior knowledge improves interpretability of chemometrics models, Chemometrics and Intelligent Laboratory Systems 240 (2023) 104905. 10.1016/j.chemolab.2023.104905. URL https://www.sciencedirect.com/science/article/pii/S0169743923001557

[14] R. Caruana, Multitask learning, Machine Learning 28 (1) (1997) 41–75. doi:10.1023/A:1007379606734.

[15] S. Ben-David, R. Schuller Borbely, A notion of task relatedness yielding provable multiple-task learning guarantees., Machine Learning 73 (3) (2008) 273–287.

[16] L. Kuncheva, A stability index for feature selection, in: Proceedings of the 25th International Multi-Conference: Artificial Intelligence and Applications, ACTA Press, Anaheim, CA, USA, 2007, pp. 390–395.

